# Automated Detection of Livestock Gastrointestinal Parasite Eggs and Cysts Using YOLOv8-Based Deep Learning

**DOI:** 10.64898/2026.08.21.746375

**Authors:** Afsan Sarwer

## Abstract

Parasitic infection is one of the common health problems of livestock in Bangladesh. Due to the country’s climate, heavy monsoon rainfall, low biosecurity in farms, and high humidity, along with the presence of suitable vector organisms, gastrointestinal parasitism remains widespread in cattle and other livestock. The standard method of diagnosis is microscopic examination of fecal samples, but this depends on manual observation, which is time-consuming and can lead to error, mainly because many parasite eggs look similar to each other and samples often contain contaminants that can be mistaken for eggs or cysts. In this study, we applied the YOLOv8 deep learning model for automated detection of parasitic eggs and cysts from microscopic images of livestock fecal samples. Images of clinical cases were collected, annotated, and used to train the model in Python, with batch size 16, auto optimizer, learning rate 0.01, momentum 0.937 and weight decay 0.0005. Training was carried out using Google Colab, and the model was evaluated using precision, recall, F1-score, mAP50 and mAP50-95. The model achieved a precision of 56%, recall of 24%, F1-score of 33.6%, mAP50 of 33%, and mAP50-95 of 22%. The relatively low recall and F1-score indicate that the model still has considerable limitation, largely due to insufficient species-specific training data and the presence of image artifacts. Underrepresentation of some parasite species, such as Trichuris spp., in the dataset also caused class imbalance, which affected the model’s ability to detect these species reliably. Despite these limitations, the study indicates that the YOLOv8 architecture has some potential for detection of parasitic eggs and cysts from microscopic images, and that further work with larger and more balanced datasets may improve performance and applicability in veterinary diagnostics.

## 1. Introduction

Parasitic infection is one of the most common health problems affecting livestock in Bangladesh. The reported prevalence of gastrointestinal parasites (GIPs) in cattle in Bangladesh ranges between 72.0% and 84.8% (Sayeed et al., 2024). This high burden of parasitism is largely attributable to the country’s geographic location, the heavy rainfall associated with the monsoon, inadequate agricultural hygiene, low biosecurity, high humidity, an open grazing system, and the presence of suitable vector organisms for helminth transmission. The range of endoparasites affecting livestock in Bangladesh is wide, including nematodes, trematodes, cestodes, and protozoa. Parasitic infections can lead to reduced growth, decreased milk and egg production, and even death if left untreated, and farmers in Bangladesh often struggle to monitor and control parasitic infestation in their herds.

Veterinary surgeons routinely diagnose endoparasitic infection through microscopic examination of fecal samples. However, this process is time-consuming and requires laboratory facilities, which are frequently limited. As a result, diagnosing a large number of cases becomes difficult, particularly in rural areas where livestock rearing is common and the grazing area is extensive, but laboratory access is scarce.

Deep learning-based image analysis has been increasingly applied in medical and biological fields to assist or partially automate tasks that traditionally depend on manual visual inspection (Shen et al., 2017). Among the object detection frameworks used for this purpose, the YOLO (You Only Look Once) family of models has drawn attention for its comparatively fast inference and reasonable accuracy. Tuncer et al. (2024) applied a YOLOv8-based model for nail capillary detection to aid early diagnosis of conditions such as scleroderma, reporting an F1-score of 0.83; the authors noted that their work was still limited by the size of the training dataset and available computational resources. Dziadosz et al. (2024) used YOLOv8 for automatic detection of microorganisms such as Arcella vulgaris in microscopic images of activated sludge, reporting an accuracy of 0.9084 and noting that YOLOv8 performed faster and more accurately than earlier YOLO versions in their setting. Liu et al. (2021) pointed out that one of the main bottlenecks in applying deep learning to microscopy image analysis is the availability of reliable ground truth annotation, since manual annotation by experts, although labor-intensive, remains the most dependable way to achieve good model accuracy.

Building on this background, the present study explored the application of the YOLOv8 architecture for automated detection of parasitic eggs and cysts in microscopic images of livestock fecal samples collected in Bangladesh. The aim was not to deliver a finished diagnostic product, but to evaluate, as a preliminary step, how far a YOLOv8-based model could go in identifying these targets under real clinical sample conditions, and to identify the practical limitations that would need to be addressed before such a tool could be used reliably in the field.

## 2. Materials and Methods

### 2.1 Image collection

Microscopic images of parasitic eggs and cysts were collected from clinical fecal samples of livestock. Images were captured under consistent lighting conditions to reduce the effect of variation in light intensity between samples, since such variation can affect the appearance of eggs, cysts, and background debris in the image.

### 2.2 Image annotation and dataset preparation

Each collected image was cropped to an appropriate size to support accurate detection. Eggs and cysts of common endoparasites present in the images were manually annotated by experts using Roboflow, a platform commonly used for preparing and managing computer vision datasets. Representative examples of manually annotated targets are shown in Figure 2. After annotation, all bounding boxes were visually inspected and corrected where necessary. The annotated images were then exported in the YOLOv8-specific dataset format. The dataset was divided into training, testing, and validation subsets in a ratio of 70%, 20%, and 10% respectively (Figure 1).

**Figure 1.**
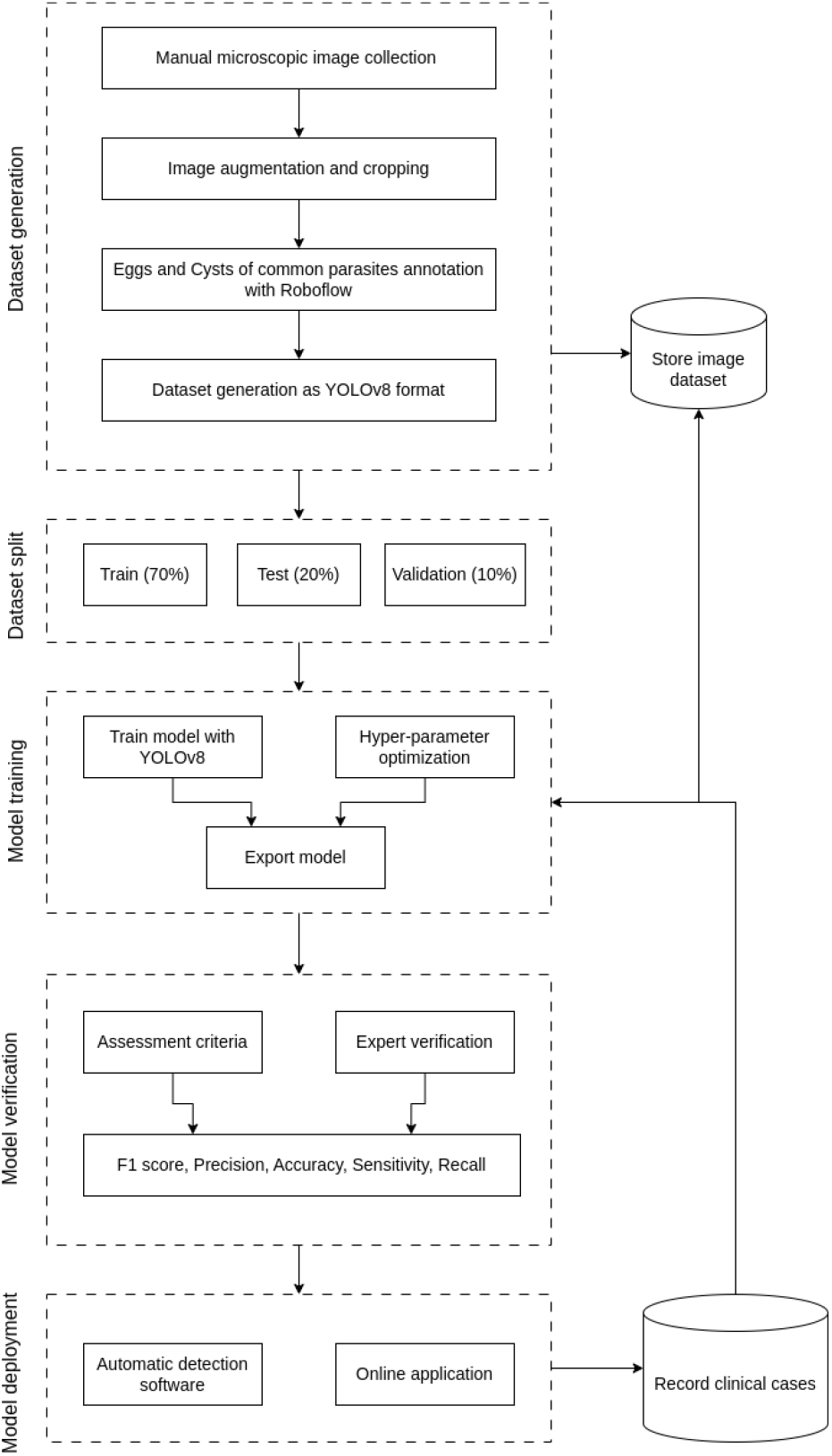
Workflow of dataset generation, model training, and evaluation used in this study.

**Figure 2.**
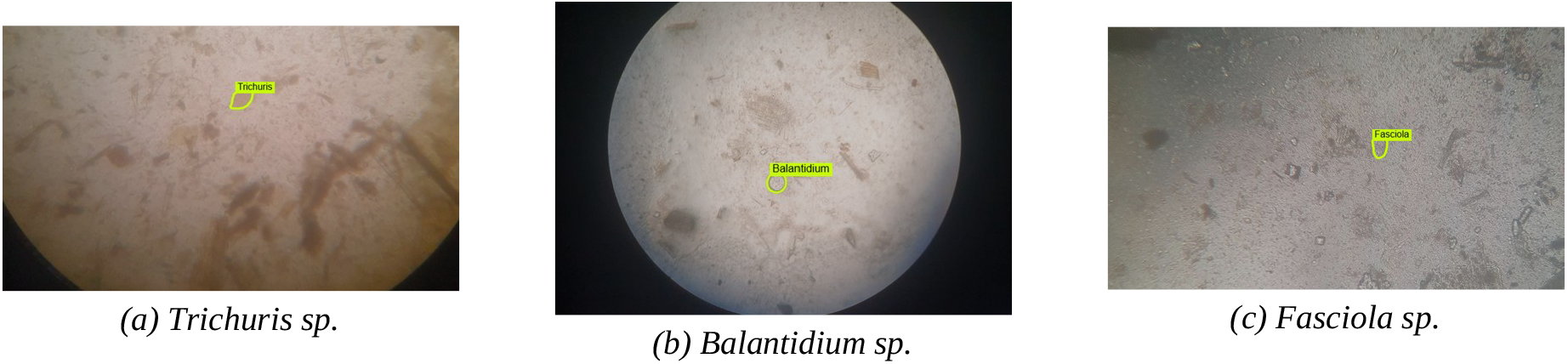
Representative examples of manual annotation of parasite eggs and cysts in microscopic images: (a) Trichuris sp., (b) Balantidium sp., (c) Fasciola sp.

### 2.3 Model training

The YOLOv8 object detection architecture (Ultralytics) was implemented using the Python programming language. YOLOv8 was selected for this study because it has been reported to offer a favorable balance of speed and accuracy compared with several earlier object detection approaches, which was considered useful given the eventual goal of near real-time diagnostic support. The model was trained on the annotated dataset using Google Colab, with the following hyperparameters: batch size of 16, auto optimizer, learning rate of 0.01, momentum of 0.937, and weight decay of 0.0005.

### 2.4 Performance evaluation

After training, the model was used to generate predictions on the held-out test image set. Model performance was assessed using a confusion matrix, in which true positives (TP) represent cases where the model correctly identified a target, true negatives (TN) represent correct identification of the absence of a target, false positives (FP) represent incorrect identification of a target that was not present, and false negatives (FN) represent a missed target that was actually present (Figure 3). Based on this matrix, the model was evaluated using precision (the proportion of correctly detected eggs/cysts among all detections made by the model), recall (the proportion of correctly detected eggs/cysts among all eggs/cysts actually present in the dataset), F1-score (the harmonic mean of precision and recall), mAP50 (mean average precision at an intersection-over-union threshold of 0.50), and mAP50-95 (mean average precision averaged over IoU thresholds from 0.50 to 0.95). Predictions were additionally checked against expert verification.

**Figure 3.**
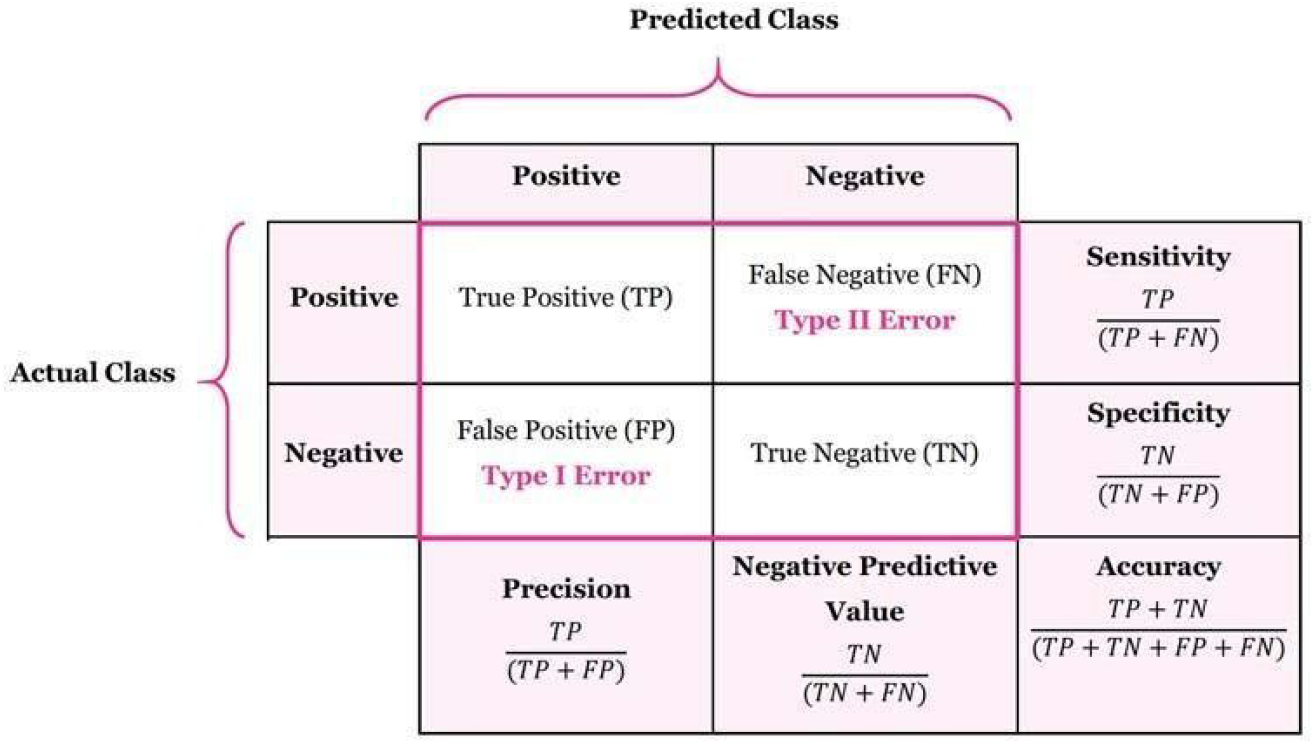
Confusion matrix and derived performance metrics used to evaluate model predictions.

## 3. Results

The YOLOv8 model, trained on the annotated dataset of microscopic images of livestock fecal samples, achieved an average precision of 56%, a recall of 24%, and an F1-score of 33.6% on the test set. The mAP50 was 33% and the mAP50-95 was 22% (Table 1). Training and validation loss decreased steadily across epochs, while precision, recall, and mAP metrics improved over the course of training before plateauing in the later epochs (Figure 4).

**Table 1.** Performance of the YOLOv8 model on the test dataset.

| Metric | Value |
| --- | --- |
| Precision | 56% |
| Recall | 24% |
| F1-score | 33.6% |
| mAP50 | 33% |
| mAP50-95 | 22% |

**Figure 4.**
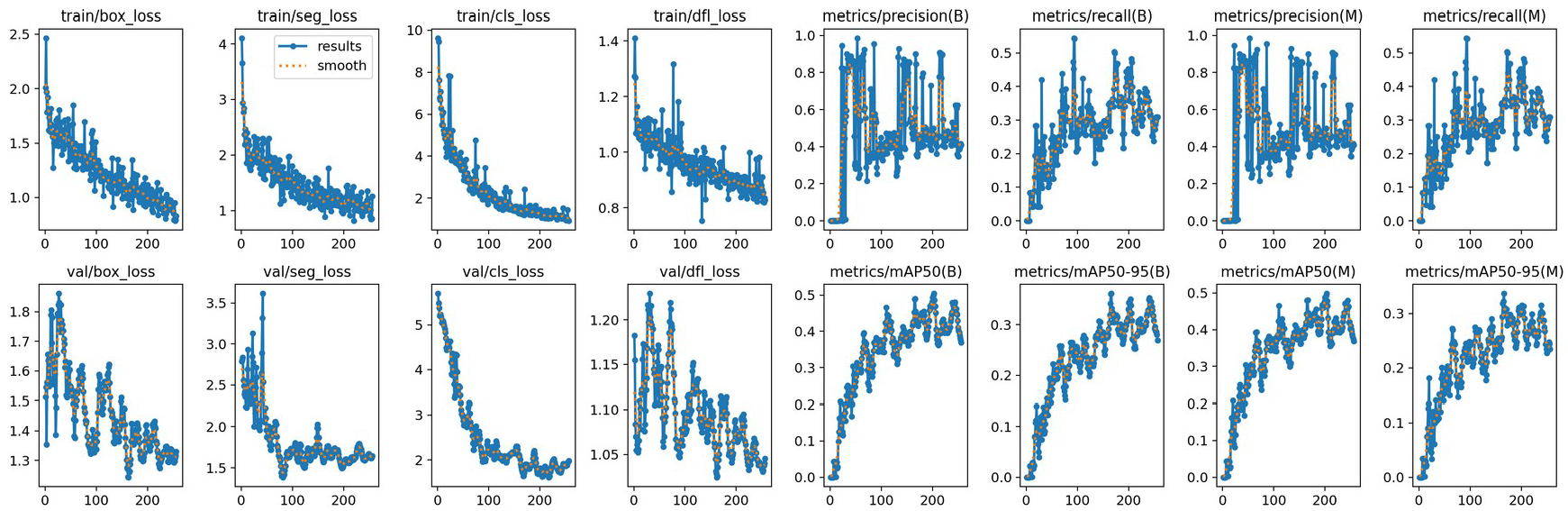
Training and validation loss and performance curves for the YOLOv8 model across training epochs (B: bounding-box metrics; M: mask/segmentation metrics).

Model performance was not uniform across all target classes. Certain parasite species, such as Trichuris spp., were underrepresented in the training dataset, and this class imbalance appeared to reduce the model’s ability to detect these species reliably compared with more frequently represented ones. In addition, the presence of image artifacts, such as debris and other contaminants commonly found in fecal samples, appeared to contribute to missed detections and, in some cases, false detections. Representative examples of instances where the model correctly detected and classified a target with reasonably high confidence are shown in Figure 5; these serve only as illustrative examples of correct output and do not reflect the average performance reported above.

**Figure 5.**
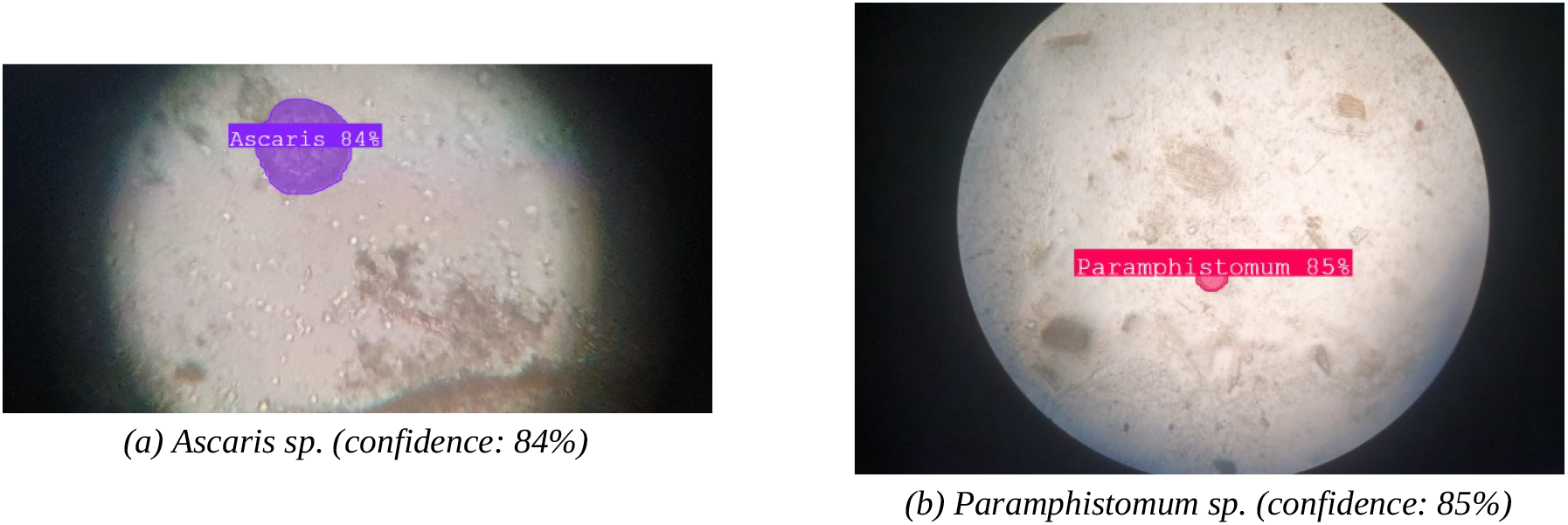
Representative examples of correct detections by the trained model, with predicted confidence scores. These illustrate individual correct outputs and are not representative of average model performance across the full test set.

## 4. Discussion

The precision obtained in this study (56%) suggests that when the model did identify a target, it was correct slightly more often than not, but the low recall (24%) indicates that the model missed a substantial proportion of eggs and cysts actually present in the images. This gap between precision and recall, reflected in the modest F1-score of 33.6%, points toward a model that is still under-trained relative to the complexity of the task, rather than one that is fundamentally unsuited to it.

This level of performance is lower than what has been reported in some related applications of YOLOv8 to microscopic or biomedical image analysis. Tuncer et al. (2024) reported an F1-score of 0.83 for YOLOv8-based nail capillary detection, and Dziadosz et al. (2024) reported an accuracy of 0.9084 for detection of microorganisms in activated sludge. Both of these studies, however, worked with comparatively controlled and homogeneous imaging conditions. In contrast, livestock fecal samples are inherently heterogeneous, containing a wide variety of debris, plant material, and other contaminants alongside the target eggs and cysts, and different parasite species can appear morphologically similar to one another under the microscope. This is broadly consistent with the observation of Shen et al. (2017) that medical image analysis models often need to contend with images acquired under varying conditions, and that handling this variability remains a practical challenge for deep learning-based approaches.

The class imbalance observed for species such as Trichuris spp. is also consistent with the point raised by Liu et al. (2021) that the quality and completeness of ground truth annotation is a central limiting factor for deep learning-based image analysis. In the present study, annotation depended on the availability of clinical cases in which particular species happened to be present, and less common species were therefore represented by fewer training examples. This is a common constraint in clinical veterinary settings, where sample collection cannot always be controlled to ensure a balanced representation of all target classes.

Taken together, these results should be read as an early, feasibility-stage finding rather than evidence of a diagnostic tool ready for field use. The dataset used in this study, drawn from a limited number of clinical cases at a single institution, was almost certainly not large or diverse enough to allow the model to learn robust, generalizable features for every parasite species of interest. Expanding the dataset, both in overall size and in the representation of less common species, together with more systematic control of image quality during sample collection, would likely be necessary before performance could approach the levels reported in more controlled applications of YOLOv8.

## 5. Conclusion

This study explored the use of the YOLOv8 deep learning architecture for detecting parasitic eggs and cysts in microscopic images of livestock fecal samples collected in Bangladesh. The model demonstrated that automated detection of this kind is feasible in principle, but its current performance, with a precision of 56% and recall of 24%, remains modest and is not yet sufficient for reliable clinical use. Insufficient species-specific training data, class imbalance, and the presence of image artifacts were identified as the main factors limiting performance. Future work with larger and more balanced datasets, covering a wider range of parasite species and sample conditions, will be needed to improve the model’s reliability and move it closer to practical application in veterinary diagnostics.

## 6. Future Work

Following the establishment of this foundational framework under controlled laboratory conditions, an industrial-grade microscopic camera, such as the Imaging Source DFK 38UX267, could be employed for in-situ identification of helminths in the field. Field-based diagnosis of this kind could help reduce diagnostic turnaround time, lower the chance of human error, and reduce dependence on centralized laboratory procedures, provided that the underlying model is first improved using a larger and more balanced dataset. A dedicated software application, built using the Python Tkinter graphical interface and linked to a microscope, is also planned to serve as a digital repository of clinical cases, supporting ongoing refinement of the model over time (Figure 6).

**Figure 6.**
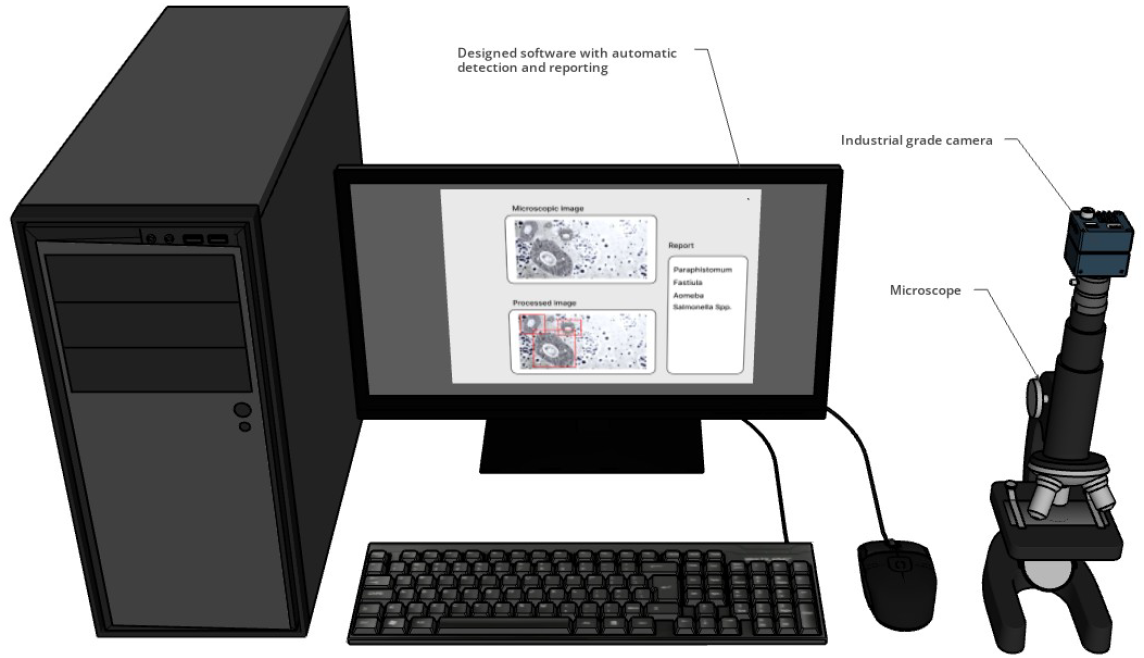
Conceptualized setup of the proposed field diagnostic system, integrating a microscope, industrial-grade camera, and detection software (planned future work).

## Acknowledgement

We would like to extend our sincere gratitude to Dr. AKM Anisur Rahman, Professor, Department of Medicine, Faculty of Veterinary Science, Bangladesh Agricultural University, for his valuable guidance and support throughout this study. His expertise and mentorship played a significant role in shaping this study.

## Notes

### Competing Interest Statement

The authors have declared no competing interest.

